# Correlated spontaneous activity sets up multi-sensory integration in the developing higher-order cortex

**DOI:** 10.1101/2024.07.19.603239

**Authors:** JaeAnn M. Dwulet, Nawal Zabouri, Jan H. Kirchner, Marina E. Wosniack, Joey R. Stawyskyj, Alessandra Raspanti, Deyue Kong, Gerrit J. Houwen, Paloma P. Maldonado, Christian Lohmann, Julijana Gjorgjieva

## Abstract

To perceive and navigate complex sensory environments, animals combine sensory information from multiple modalities in specialized brain circuits. Known as multisensory integration, this process typically depends on the existence of co-aligned topographic connections from several sensory areas to downstream circuits exhibiting multimodal representations. How such topographically co-aligned connectivity necessary for multisensory integration gets set up in early stages of development is still unknown. Inspired by the role of spontaneous activity in refining topographic connectivity between early sensory circuits, here we investigated the potential of such spontaneous activity to also guide the co-alignment of multiple sensory modalities in RL, a higher-order associative cortical area rostro-lateral to V1. Analyzing spontaneous activity simultaneously recorded in primary visual and somatosensory cortex and area RL at different developmental ages before sensory experience, we identify candidate features of this activity to guide the emergence of co-aligned topographic multisensory projections with somatosensory leading the visual projection. We confirm this hypothesis using a computational model of activity-dependent circuit refinement, and show that the correlation of spontaneous activity between the visual and somatosensory primary cortex can establish an optimal fraction of multisensory neurons in RL for stimulus decoding. Our model provides an exciting new computational perspective of the role of spontaneous activity in the emergence of topographically co-aligned multimodal sensory representations in downstream circuits, specialized for the processing of rich sensory environments.

## Introduction

Sensory environments are complex – rich with objects and events that specific brain circuits must decipher to ensure their accurate perception and to guide appropriate behavior. To efficiently perceive and navigate sensory environments, these brain circuits combine information from multiple sensory modalities, including vision, somatosensation, or hearing, a process known as multisensory integration. Several brain regions have been shown to generate multimodal representations, typically by aligning connections from the sensory periphery where specific sensory features are first encoded to downstream projection areas. In the rodent, whiskers transmit tactile sensations which are combined with visual information to guide many behaviors including goal-directed behavior [1], gap crossing and food source detection [2], navigation [3], prey-capture behavior [4], and recognizing object features and shapes [5]. Multisensory integration has been reported in subcortical areas such as the superior colliculus [6, 7] and the striatum [8]. Higher-order areas (HOAs) also play play a critical role in multisensory integration, because they receive and combine inputs from multiple primary sensory areas [1, 3, 9–11]. Some of these higher-order areas, including RL, are anatomically positioned between the primary sensory cortices whose inputs they integrate. Moreover, connectivity is organized in a topographic manner across the modalities, whereby neurons representing neighboring locations in sensory space project to neighboring locations in higher-order cortex [3, 12]. For multisensory integration, this means that visual and somatosensory inputs representing corresponding regions of space can converge onto overlapping or nearby locations in higher-order cortex. Specifically, visual and tactile stimuli have overlapping representations in a higher order association area, area RL, that lies between the primary visual (V1) and the primary somatosensory (S1) cortices and receives topographically organized input from both areas. There is significant spatial alignment between V1 and S1 in area RL; e.g. the lower whisker pad aligns to the lower visual field of view and can elicit responses in the same neurons, called bimodal neurons, in RL [3]. RL is therefore a particularly useful area for studying how topographically aligned visual and somatosensory inputs are integrated in higher-order cortex and how such aligned multisensory representations emerge during development. Additionally, this multisensory integration allows for cross-modal generalization. For example, rodents can adapt to changing sensory environments and can generalize sensorimotor tasks to a different modality, i.e. visual, after learning a task from only tactile stimulus, due to aligned abstract representations of these modalities in RL [1], further supporting RL as a relevant model system for studying multi-sensory integration. Finally, evidence shows that rats combine and integrate visual and tactile signals in a supralinear manner, significantly enhancing sensory processing efficiency [5]. This integration is supported by a substantial population of bimodal neurons in RL, which respond to both visual and tactile inputs. Although full functional maturation of visual HOAs occurs only after eye opening, basic properties such as responsiveness to stimuli and topographic organization between V1 (as well as S1) and RL are already present at eye opening [13, 14]. These anatomical and functional properties make RL an ideal model system for asking how aligned multisensory representations are established before sensory experience.

Before the onset of sensory experience, primary sensory cortices generate spontaneous activity, which plays a pivotal role in their proper maturation in terms of cellular properties and activity-dependent connectivity refinements [15–18]. Across the sensory areas, a general developmental profile of spontaneous activity can be observed: early global and dense spontaneous events transition to more localized and sparse patterns, typical of adult networks [15, 17, 19–21]. By promoting the refinement of appropriate connectivity [22–24], these spontaneous activity patterns prepare the developing cortex for future sensory input [25–27], but at the same time are affected by changing circuitry [28]. Spontaneous activity has been characterized and shown to guide the establishment of hierarchical architectures in HOAs in the visual system before eye opening [29], but much less is known about its role in the development and topographic co-alignment of multiple sensory modalities in multisensory HOAs.

Whereas activity-independent mechanisms are known to set up the first cues instructing axons where to go [30, 31], activity-dependent synaptic plasticity mechanisms instructed by either spontaneous activity or sensory experience, are crucial in refining this imprecise connectivity [22, 32, 33]. Indeed, various forms of Hebbian synaptic plasticity, which promotes the potentiation of synaptic strength as a result of correlated pre- and postsynaptic activity, have been proposed to guide the activity-dependent refinement of circuit connectivity in development [28, 34–38]. Some experimental work has also measured Hebbian-like learning rules specific to development [39]. Activity-dependent mechanisms likely also contribute to the establishment and alignment of maps from different sensory modalities to downstream circuits, which has been mainly investigated in the rodent superior colliculus and avian optic tectum [32]. For example, how visual experience aligns the auditory and visual maps during development has been mainly studied by manipulations that cause the two maps to misalign [40–44]. This suggests that these same principles might apply not only to realign and maintain maps after manipulation, but also to the activity-dependent emergence of aligned representations in developing circuits.

Here, we combine computational modeling and calcium imaging experiments to investigate the role of spontaneous activity in the emergence of aligned topographic maps from two primary sensory cortices (visual, V1 and somatosensory, S1) to RL, an associative cortical area rostro-lateral to V1, and the emergence of bimodal responses in RL. We find that spontaneous activity in V1, S1, and RL sparsifies during development, but S1 exhibits more mature activity patterns compared to V1, which desynchronize as early as postnatal day eight. Using these experimental insights, we expand a computational model of activity-dependent connectivity refinement between primary sensory areas (V1 and S1) and RL using biologically plausible Hebbian synaptic plasticity which was previously used to study topographic map-formation and receptive field refinement between the thalamus and V1 [28]. Our model predicts that aligned topographic connectivity refinement and the emergence of bimodal neurons requires an intermediate amount of correlation between the primary sensory cortices (V1 and S1), which we investigate in our data. This connectivity refinement through spontaneous activity between the primary sensory areas and RL is sufficient to explain the developmental sparsification in RL whereby RL neurons become responsive to either or both primary sensory cortices in a ratio that ensures optimal information processing.

## Results

### Spontaneous activity in V1, S1 and RL during development

To investigate how spontaneous activity matures differentially across the mouse’s developing primary visual (V1), somatosensory (S1), and higher-order areas, specifically area RL, we performed two-photon imaging with one field of view per developing cortex in unanesthetized mouse pups at postnatal days (PN) 8-16 (Figure 1A,B). Spontaneous activity was observed across all three cortical areas (Figure 1C). We used a linear mixed model analysis [45] to estimate the differential effect of age on different aspects of spontaneous activity. Consistent with previous studies of developing visual cortex [15, 17, 21, 46], we found that spontaneous activity in V1 sparsifies throughout development, i.e. the average amplitude, duration, and participation rate of spontaneous activity decrease with age, as reflected in the significant negative effect of age on these properties (Figure 1D-F, H, see Methods). In contrast, the frequency of events increases with age, as reflected in the significant positive relationship between age and event rate (Figure 1G, H). Similar trends of activity sparsification were also observed in S1 and RL. Thus, while the developmental sparsification of spontaneous activity corroborates previous work in primary sensory cortex, our analysis extends these observations by comparing V1, S1, and RL within the same experimental and statistical framework. There were no differences between participation rate across areas, with all areas following a similar decreasing trend over time (Figure 1F,H). Furthermore, comparing area RL to V1 also showed no significant differences in duration of events and event rate between these two areas (Figure 1E,G,H), although amplitudes were slightly higher in RL compared with V1. However, we observed some significant differences, especially in S1. Specifically, spontaneous events in S1 have significantly lower amplitudes, lower durations, and higher rates compared to the other regions, at younger ages (Figure 1D,E,G,H).

**Figure 1.**
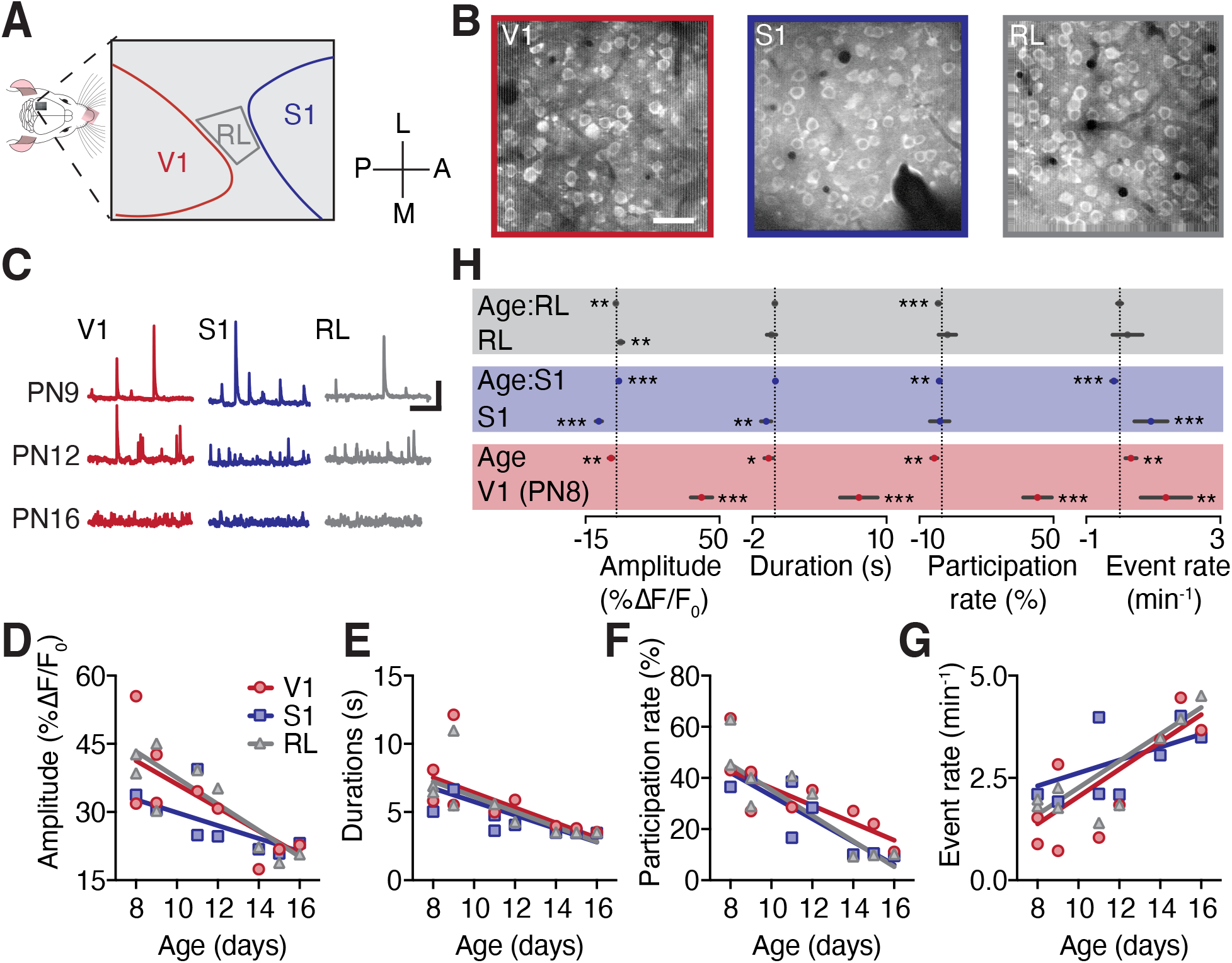
Spontaneous activity in RL sparsifies over development, with more mature activity in S1 compared to V1 and RL.**A.**Schematic of the two-photon experiments. Primary visual cortex (V1), somatosensory cortex (S1), and rostro-lateral area (RL) imaged in sequence over a 100 min recording session. **B**. Sample field of views of the individual cortices. Scale bar corresponds to 50 *µ*m. **C**. Sample z-profile traces from the three different cortices (columns) in three mice at different postnatal ages (rows). Vertical scale bar corresponds to 20% ΔF*/*F_0_. Horizontal scale bar corresponds to 50 seconds. **D-G**. Amplitude (D), duration (E), participation rate (F), and event rate (G) as a function of postnatal age. Each dot represents the average value for one animal for one cortical area at the indicated postnatal age, computed across all detected events in that area. *N* = 9 was the total number of animals included across all ages. Because animals were distributed across postnatal ages and some points overlap visually, fewer than 9 points are visible at any individual age. Regression lines derived from the linear mixed model in H. **H**. Linear mixed-model estimates for each activity feature, where age and region are included as fixed effects, and mouse ID and sample ID are included as random effects. Values are shown relative to V1 at PN8. The V1 term corresponds to the model intercept, the Age term corresponds to the age-dependent slope in V1, and the S1, RL, Age:S1, and Age:RL terms indicate differences from this reference (see Methods). Stars indicate statistical significance (* p*<*0.05, ** p*<*0.01, *** p*<*0.001). Because intercepts and slopes have different units and scales, full model estimates, confidence intervals, and comparisons using S1 or RL as reference areas are provided in Supplementary Tables S1-S3.

Together, this suggests that spontaneous activity in S1 has more mature amplitudes, duration, and event rates earlier in development, whereas RL and V1 develop more similarly during the second postnatal week. In summary, spontaneous activity in developing primary cortices and RL sparsifies with different temporal profiles, with the somatosensory cortex showing earlier maturation than the visual cortex and RL (Supplementary Tables S1-S3).

### The primary somatosensory and visual cortex exhibit correlated spontaneous activity

Having established that spontaneous activity matures with different temporal profiles across V1, S1, and RL, we next asked whether activity in V1 and S1 is temporally and spatially correlated in a way that could support the emergence of aligned multisensory representations in RL. To this end, we reanalyzed our published wide-field calcium imaging recordings from both V1 and S1 [47] to assess the temporal relationship between spontaneously occurring activity in these two primary sensory cortices (Figure 2A). We found that activity between the visual and the somatosensory cortex is often, but not always, temporally synchronized, with examples showing near-synchronous but spatially distinct activation of subregions in V1, RL, and S1 (Figure 2B-D). Indeed, analyzing the average activity between V1 and S1 of several animals at postnatal days PN9-PN12 revealed a range of Pearson correlation coefficients, with a mean of ~0.5 (Figure 2H).

**Figure 2.**
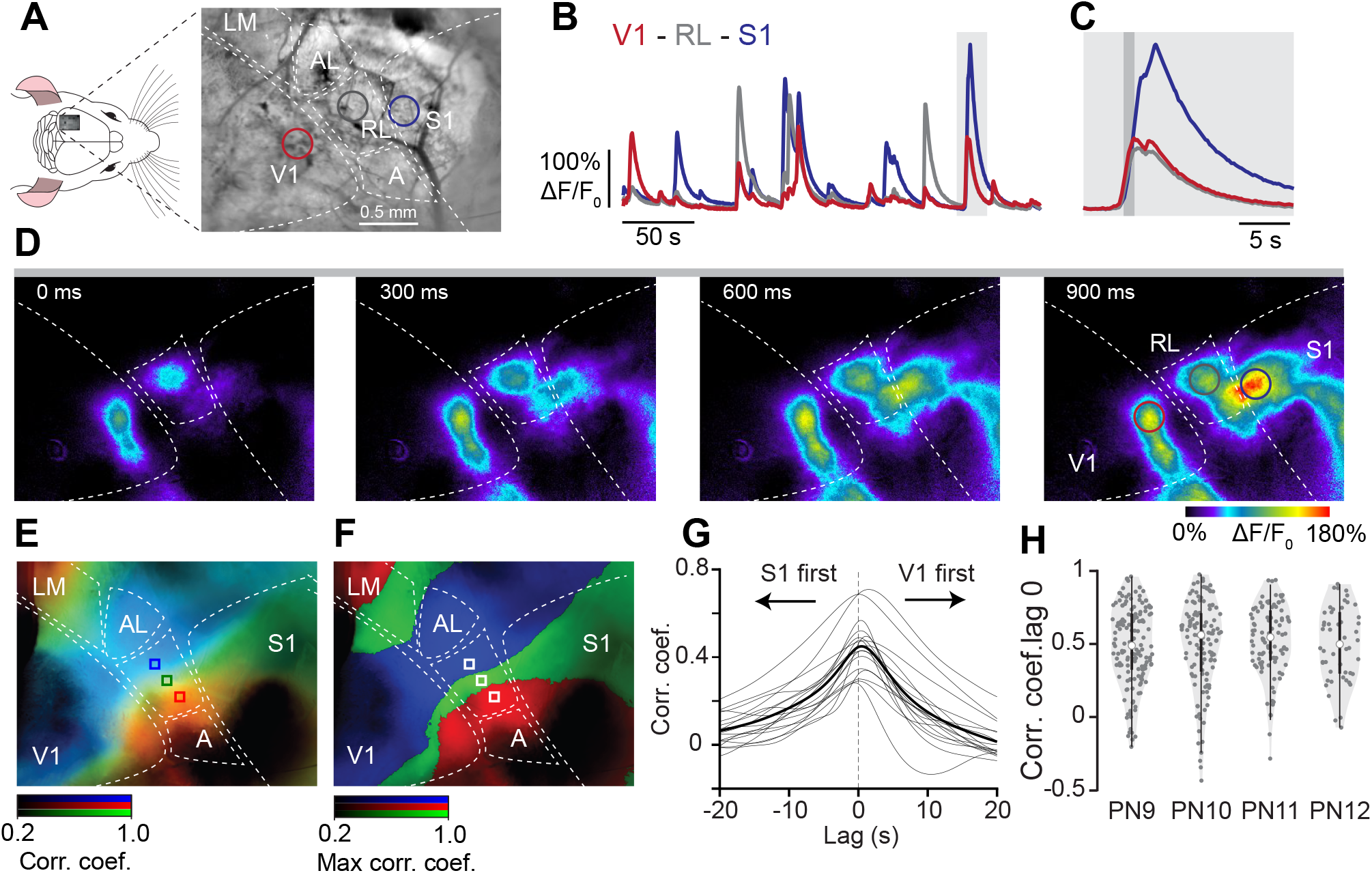
Spontaneous activity in V1, S1, and RL is near-synchronous and spatially organized. **A.**Schematic of the wide-field experiments. Cortical surface area encompassing the primary visual (V1) and somatosensory (S1) cortices as well as RL in a PN9 mouse. Higher-order areas of the visual cortex are drawn according to [9]. LM: lateromedial area, AL: anterolateral area, A: anterior area. **B**. ΔF*/*F_0_ traces from 3 ROIs in V1, S1 and RL (shown in A). **C**. Expanded view of the traces shown in the area marked by a light grey box in B. **D**. Four ΔF*/*F_0_ frames over a period of 900 ms indicated by a dark grey vertical bar in C. **E**. Functional correlation map of the recording shown in B-D. Pixel values represent Pearson correlation coefficients between each pixel and the areas marked by blue, green and red squares in RL across the recording’s duration. The color of each pixel is the RGB composite of the coefficients of the areas marked by the three colored squares in RL. **F**. Same as E except that each pixel is assigned to the color channel (red, green or blue) with the highest coefficient of correlation value across the three channels. **G**. Pearson correlation between activity in somatosensory and visual cortex as a function of temporal lag at PN9. Individual animals are shown in gray and the mean in black. The vertical black line indicates zero lag. Arrows indicate the temporal order of activity at different lags. **H**. Temporal correlations between V1 and S1. Violin plots where each dot indicates the Pearson correlation coefficient for one 2-minute bin. PN9 *N* = 6; PN10 *N* = 5; PN11 *N* = 4; PN12 *N* = 2.

To test whether these correlations reflected a consistent propagation of activity from one primary sensory cortex to the other, we computed V1–S1 correlations at PN9 as a function of temporal lag. Correlations were maximal near zero lag and decreased for both positive and negative lags, arguing against stereotyped traveling-wave-like propagation between V1 and S1 (Figure 2G). The lagged correlation curves showed a mild asymmetry, with somewhat stronger correlations when S1 preceded V1. However, because the dominant peak was near zero lag, we interpret these data primarily as evidence for near-synchronous coactivity rather than a consistent temporal lead of one area over the other. Together with the examples of near-synchronous but spatially distinct activation in V1, RL, and S1, this suggests that spontaneous activity across these areas is predominantly near-synchronous and spatially structured, rather than reflecting a single wave propagating across the entire imaged region.

Previous work has shown that connectivity between V1 and RL as well as S1 and RL is topographically organized in mature animals to preserve space [3, 9]. Furthermore, this organization is also aligned across the modalities, i.e. stimuli in the same region of sensory space tend to converge onto the same cells in

RL [3, 12, 29]. Hence, in addition to temporal correlations, we evaluated whether spontaneous activity also carries information about spatial topography between the two primary sensory cortices and RL. We computed functional correlation maps for three distinct seed locations in RL, showing that spontaneous activity in RL is coordinated topographically with activity in both V1 and S1 (Figure 2E,F). Additional functional-correlation-map examples from PN9, PN10, and PN13 recordings showed similar spatially ordered correlation structure across the developmental window analyzed here, with seed locations placed in V1, S1, or RL as indicated in each panel (Supplementary Figure S1). This is consistent with previous work showing that functional-connectivity maps computed from spontaneous activity can reveal topographic organization in higher visual areas before eye opening [29], and with recent evidence that retinotopy-like and somatotopy-like patterns of ongoing activity, together with their rough topographic correspondence in RL, are already present at P10–11 [48]. Thus, spontaneous activity between V1 and RL and between S1 and RL is also spatially correlated and carries topographic information. This suggests that some activity-dependent mechanism might be using these temporal and spatial correlations during development to establish topographic connectivity maps between the primary sensory areas and RL, and hence enable multisensory integration in the adult animal.

### Correlated spontaneous activity can establish topographic connectivity between the primary sensory cortex and RL and generate bimodal cells in RL

To test the hypothesis that an activity-dependent mechanism may use the spatiotemporally structured spontaneous activity in the primary sensory cortices to set up topographic connectivity from each area into RL and generate bimodal cells, we built and analyzed a computational model with three populations of neurons for V1, S1, and RL (Figure 3A) [28]. We modeled the activity-dependent synaptic plasticity of feedforward connections between V1 and RL, and S1 and RL, with each area subject to spontaneous activity as characterized before (Figure 1). While we initialized the connectivity as random, we included a slight bias for neighboring cells in the primary sensory cortices to project onto neighboring cells in RL (Figure 3B, Methods). This bias incorporates activity-independent effects like molecular gradients that are known to set up a coarse topography before the onset of activity-dependent plasticity, at least for connectivity between the sensory periphery and subcortical or primary cortical areas [38, 49–51]. We therefore interpret this bias as a coarse initial scaffold on which activity-dependent plasticity acts, rather than as a mechanism that by itself determines the final topographic organization. Activity-dependent plasticity was implemented through a synaptic plasticity rule involving Hebbian and heterosynaptic terms. The Hebbian term potentiates synaptic weights based on coincident pre- and postsynaptic activity. The additional heterosynaptic term was implemented to depress synaptic weights, but only in the presence of presynaptic activity (see Methods)

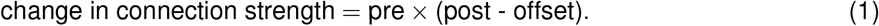

**Figure 3.**
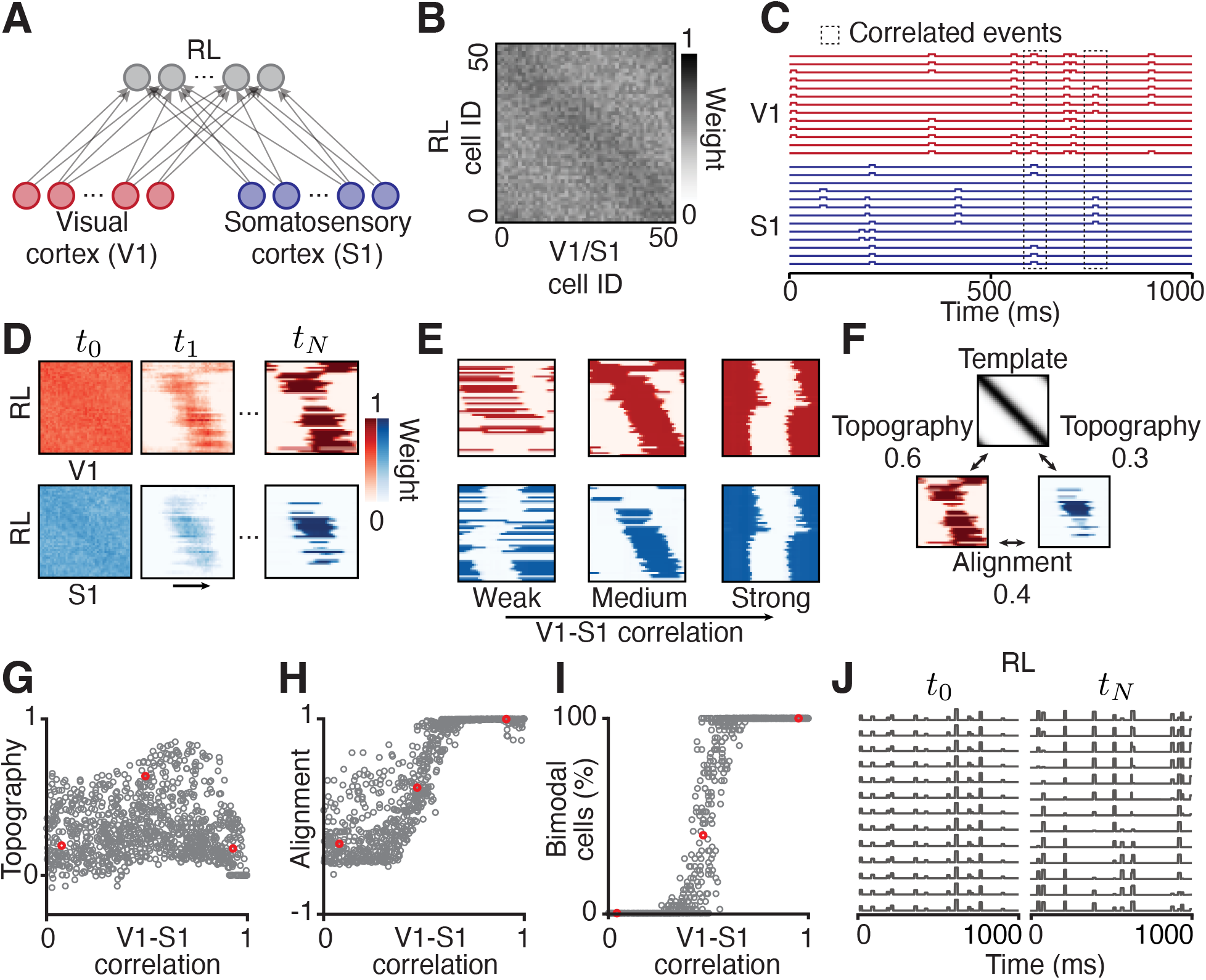
Model set-up and assumptions. **A.**Schematic of a three-population feedforward model of the developing V1, S1, and RL. **B**. Example of random initial connectivity matrix between V1 and RL (connectivity matrix for S1 and RL is chosen identically) with a topographic bias denoted by stronger connections along the diagonal inspired by activity-independent mechanisms (see Methods). **C**. Sample of spontaneous events in V1 (top) and S1 (bottom) across cells as a function of time. Boxes highlight two spatiotemporally correlated events in V1 and S1 (see Methods). **D**. Example of refinement of connectivity matrix between V1 and RL (top) and S1 and RL (bottom), from *t*_0_ to *t*_*N*_. Same axes as in Figure 2B. **E**. Example of connectivity matrices in steady state from simulations with low (correlation=0.1, left), medium (correlation=0.5, center) to highly (correlation=0.9, right) temporally correlated activity between V1 and S1. **F**. Schematic to illustrate topography and alignment measures (see Methods). **G-I**. Topography (G), alignment (H), and percentage of bimodal cells (I) as a function of V1 to S1 correlation. Red dots correspond to the examples from E. Each point represents an independent simulation run in which both the initial connectivity matrix and the stochastic sequence of spontaneous events were resampled to mimic different developmental realizations. **J**. Excerpt of activity in RL at the beginning (left) and end (right) of the simulation (corresponds to panel D, left and right).

This phenomenological plasticity rule has a biophysical basis and was previously derived from a set of interacting molecules (in particular, neurotrophins) [39, 52, 53] and shown to produce selectivity to correlated inputs from random initial connectivity [37, 54]. We then implemented spatiotemporally correlated spontaneous activity in areas V1 and S1, as measured experimentally, which this synaptic plasticity rule used to modify the synaptic weights in the network (Figure 3C). Specifically, S1 and V1 neurons were driven by a combination of independent events within each area and shared events across the two areas. Independent events activated randomly chosen contiguous groups of neurons within either V1 or S1, implementing spatial correlations within each area, which we have previously shown can establish topographic connectivity between two areas [28]. Shared events implemented spatiotemporally structured correlations between V1 and S1: they occurred simultaneously in both areas and activated the same contiguous group of neurons in terms of topographic identity. The common inputs giving rise to shared events might reflect shared inputs from the subcortical areas such as the thalamus or superior colliculus into the primary sensory cortices [55–57] or correlation induced by direct connections between V1 and S1 [58, 59].

Using this spatiotemporally correlated spontaneous activity in the primary sensory cortices, the Hebbian plasticity rule (Eq. 1) refined and stabilized connectivity between V1/S1 and RL (Figure 3D), as shown before in a two-layer network [28]. We investigated the emergent connectivity properties for a range of V1 and S1 temporal correlations (Figure 3E). To quantify the topographic organization of connectivity between V1 and RL, and S1 and RL, and the alignment of maps from V1/S1 to RL, we defined two mathematical measures, topography and alignment (Figure 3F, see Methods). Topography was defined as how well the initial topographic bias was preserved in the final receptive field between either V1/RL or S1/RL and was calculated as the correlation between the final receptive field of each sensory area with RL and an ideal receptive field (see Methods). Alignment was defined as the measure of similarity between the final receptive fields of V1 and S1 and was calculated as the correlation between them. Weak temporal correlations (where independent events dominate over shared events) resulted in strong competition between the primary sensory cortices with dominant synaptic weight depression. The strong competition generated sparse connectivity matrices to RL, with many RL neurons decoupling from the primary sensory cortices (Figure 3E, left). This resulted in poor topography and map alignment of the maps between V1/S1 and RL (Figure 3G,H). RL had almost no bimodal cells, i.e. neurons that receive input from both primary sensory cortices, in contrast to what is found in mature RL experimentally [3] (Figure 3I). Increasing temporal correlations between the primary sensory cortices increased alignment of the V1/S1 to RL maps (Figure 3H). However, very strong correlations were detrimental to V1/RL and S1/RL topography (Figure 3E,G). Strong temporal correlation between V1 and S1 rendered many RL neurons selective to visual and somatosensory inputs at the same topographic location as reflected in the identical connectivity matrices (Figure 3E, right), overriding the initial topographic bias (Figure 3B). Only a medium-strength temporal correlation between V1 and S1, as found experimentally (Figure 2C), generated a connectivity profile which prevented RL cells from decoupling from V1 and S1, while preserving topography (Figure 3E, middle).

We quantified the percentage of bimodal neurons in our model as a function of the temporal correlation between V1 and S1 (Figure 3I). By performing a steady-state analysis of our model (see Methods), we derived the critical amount of temporal correlation, *c*∗, between S1 and V1 required for the emergence of bimodal neurons. This parameter depends solely on two parameters of the model, i.e. the average firing rate of cells in the sensory cortices, ⟨pre⟩, and the heterosynaptic offset from Eq. 1,

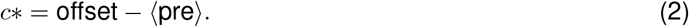

Together, these results demonstrate that the initial topographic bias was not sufficient to determine the final map structure. Instead, appropriate topography, alignment, and bimodality emerged only for an intermediate level of V1-S1 correlation (Figure 3B,E,G-I).

In a representative simulation, the refinement of V1/S1-to-RL connectivity was accompanied by sparser RL activity, visible as reduced event amplitude and participation rate over the course of learning (Figure 3J). This was consistent with the sparsification observed in the experimental data (Figure 1C,D,F) as well as with the activity sparsification previously quantified in the two-layer model of developmental refinements [28].

In summary, our proposed computational model shows how correlated spontaneous activity between V1 and S1 can refine initially biased feedforward connectivity to RL. The model predicts that only medium-strength correlations between primary cortices produce topographically organized and aligned connectivity in V1/RL and S1/RL, and a mixture of bimodal and unimodal neurons in RL consistent with experimental data.

### More mature S1 to RL connectivity can contribute to map alignment between V1 and RL

Having established that the spatiotemporal correlations between V1 and S1 can set up topographic and aligned connectivity to RL as well as generate bimodal cells in RL, we next turned to investigate a possible role of the differentially more mature spontaneous activity in S1 compared to V1 (Figure 1H). To examine this, we systematically varied the bias of the topographic connectivity between S1 and RL (Figure 4A). While a stronger bias improved the topographic connectivity between S1 and RL (Figure 4B, right), it also improved the topographic organization of connectivity between V1 and RL (Figure 4B, left, and C) as well as the alignment of connectivity from V1 and S1 to RL (Figure 4D) and affected the number of bimodal cells in RL (Figure 4E). Significantly reducing the connectivity bias from S1 to RL while using more mature S1 activity properties, including lower event amplitude and higher event frequency as observed experimentally (Figure 1D-G), quickly deteriorated topography and alignment (Supplementary Figure S2). Thus, more mature S1-like activity patterns alone were not sufficient to generate appropriately topographic and aligned maps. Rather, an initial S1-to-RL connectivity bias was required, together with the appropriate level and structure of correlated V1-S1 activity described above (Figure 3B,E,G-I).Therefore, a more mature S1 to RL connectivity bias, together with structured spontaneous activity, may contribute to the refinement of connectivity between V1 and RL.

**Figure 4.**
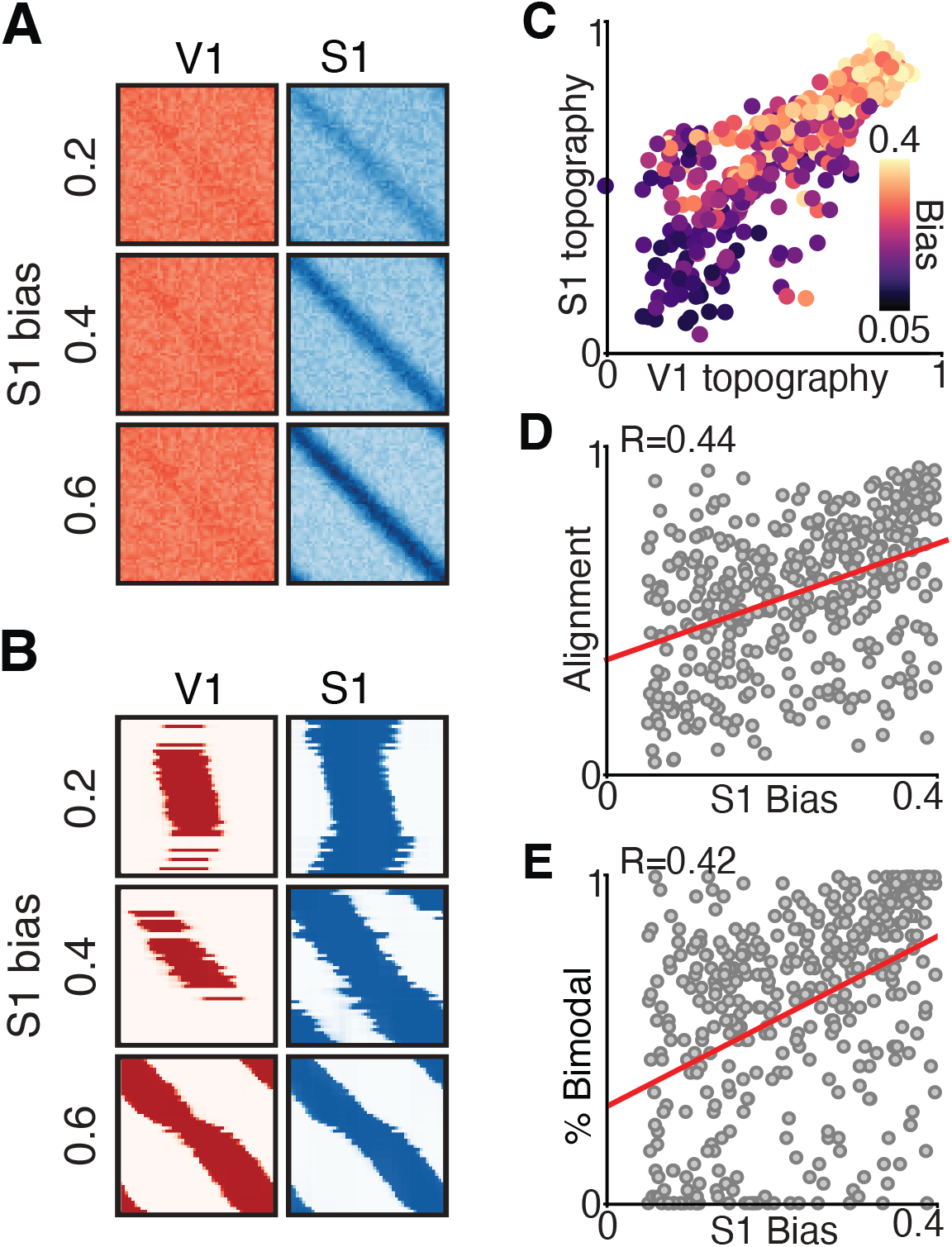
More mature spontaneous events in S1 effectively instruct map alignment between V1 and RL. **A.**Three pairs of connectivity matrices (V1 left, S1 right) with increasing (top to bottom) S1 bias. **B**. Corresponding connectivity matrices in steady-state after plasticity. **C-E**. For all simulations, V1-S1 correlation of 0.5 was used. **C** S1 and V1 to RL topography. Different S1-to-RL connectivity biases indicated by color. **D**. Alignment of V1 and S1 to RL topography as a function of S1-to-RL connectivity bias. Linear fit with *R* = 0.44, p value*<* 0.001. **E**. Percentage of bimodal neurons in RL as a function of S1-to-RL connectivity bias. Linear fit with *R* = 0.42, p value*<* 0.001.

### A mixture of bimodal and unimodal neurons in RL performs optimal decoding of activity

RL is involved in integrating information from different sensory modalities to obtain a more accurate representation of statistically rich sensory environments. In particular, an area that integrates visual and tactile stimuli is able to more accurately infer such information; e.g. when rats use both tactile and visual information, their performance in a task involving determining object orientation is better than the sum of the performance of each modality individually [5]. To investigate how the topographically aligned cortical connectivity between V1 and RL, and S1 and RL, might facilitate the integration of information from different sensory modalities, we quantified how much information about the activity of each primary sensory cortex can be decoded from the activity in RL. We quantified the amount of decoded information as the coefficient of determination, *R*^2^, which can be interpreted as the fraction of variance in V1 or S1 that can be reconstructed from activity in RL. To do this, we fitted a linear regression model using RL activity and both V1 and S1 activity, and calculated *R*^2^ from the error between our fitted linear model and each sensory area individually, which we refer to as the ‘V1 or S1 reconstruction’ (Figure 5A). [60, 61] (See Methods). *R*^2^ is naturally affected by how much activity in V1 or S1 is propagated into RL. In our model (Figure 3) and in biology [9, 62], some regions of the primary sensory cortices are not connected to RL; for example, RL contains only a partial representation of the visual field, hence V1 cells outside of that region do not project to RL [11, 63]. Consistently, we found that the fraction of variance reconstructed, *R*^2^, for either V1 or S1 independently increased with the percentage of connected neurons from the respective cortex into RL (Figure 5B, see Methods).

**Figure 5.**
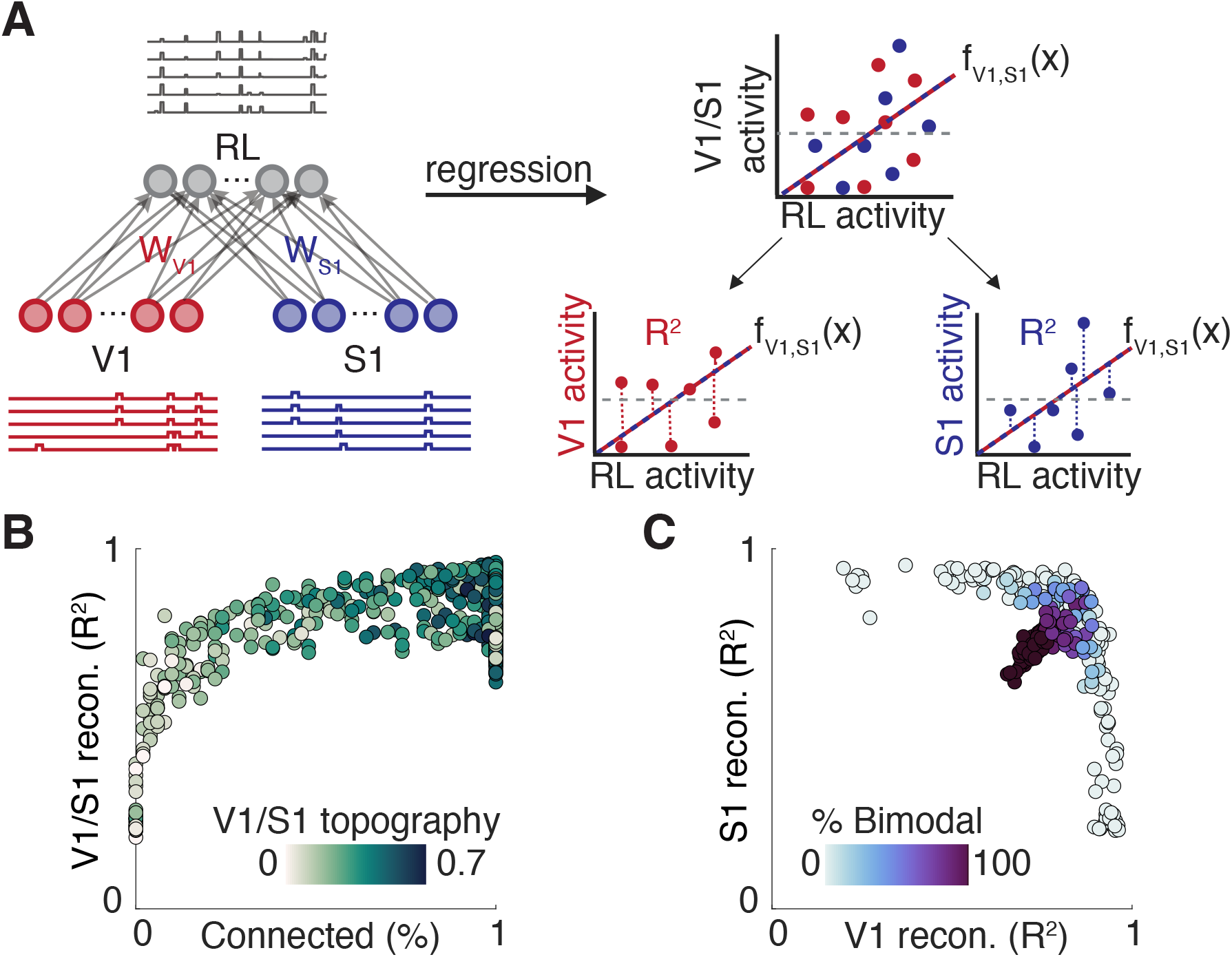
A mixture of bimodal and unimodal neurons can optimally decode activity. **A.**Schematic of the regression procedure to calculate *R*^2^ for reconstructing V1 or S1 activity from RL activity with a non-linear transfer function (a sigmoid) (see Methods). A linear regression is performed using the non-linearly transformed RL activity (dependent variable) and both V1 and S1 activity (independent variables). *R*^2^ is calculated from V1 or S1 activity separately using the fitted regression line. **B**. Reconstruction of V1 or S1 activity (*R*^2^) as a function of the percentage of connected cells from the respective cortex over all values of correlation chosen uniformly between 0 and 1 (see Methods). Color indicates topography. **C**. Reconstruction of V1 and S1 activity (*R*^2^) as a function of the percentage of bimodal cells (denoted by color). For all simulations, V1-S1 correlation of 0.5 was used.

We showed analytically that a locally connected network, where every primary sensory cortical neuron makes at least one connection to RL and every RL neuron receives at least one connection from the primary sensory cortices, can maximize the total fraction of variance reconstructed for both V1 and S1 if the activity in V1 or S1 is perfectly temporally correlated (correlation value of 1) (see Methods). These conditions correspond to a network in which all neurons in RL are bimodal. However, when activity across V1 and S1 is moderately correlated (correlation value of 0.5), this fully bimodal solution is no longer necessarily optimal. For simulations with V1–S1 moderate correlation, networks containing a mixture of unimodal and bimodal RL neurons often achieved higher joint reconstruction of V1 and S1 activity than networks composed almost entirely of bimodal neurons (Figure 5C). This occurs because unimodal RL neurons provide unambiguous information about activity in one sensory area when the two inputs are not perfectly correlated, whereas purely bimodal neurons make it more difficult to distinguish whether one or both sensory cortices were active. Thus, for moderately correlated inputs, unimodal neurons can complement bimodal neurons and improve reconstruction of both sensory areas. This suggests that the mixture of unimodal and bimodal neurons in RL [3] is the optimal solution for encoding the maximum amount of information from the activity in V1 and S1 [64].

## Discussion

We investigated the role of spontaneous activity in the maturation of connectivity from two primary sensory cortices, V1 and S1, to RL, a higher-order association area, by combining computational modeling with experiments. We found that while spontaneous activity sparsifies during development in all investigated areas, S1 exhibits relatively mature, desynchronized activity patterns as early as postnatal day eight. Using these experimental insights, we constructed a computational model of activity-dependent connectivity refinement from each V1 and S1 to RL. To generate topographic connectivity from V1 and S1 to RL, aligned connectivity maps for the two sensory projections and a mixture of unimodal and bimodal neurons in RL, our model predicts that spontaneous activity between V1 and S1 should exhibit moderate temporal correlations, which we also found experimentally. Furthermore, we demonstrated how a more mature somatosensory cortex can guide the refinement of visual cortex connectivity to RL, suggesting that inter-cortical influence plays a central role in the establishment of refined and topographically aligned connectivity during development. Finally, we found that moderate correlations between V1 and S1 result in a mixture of bimodal and unimodal neurons in RL which can optimize information processing in RL when either one modality or both are active. Thus, by combining modeling with experiments, we found that activity-dependent plasticity can instruct the formation of refined and topographically aligned connectivity between primary sensory and high-order cortices in development to produce efficient information integration from multiple modalities.

### Origin of correlated events in the primary sensory cortices

We find that early spontaneous activity with medium correlations can lead to the optimal amount of bimodal neurons and alignment of these sensory maps before sensory experience, which is crucial for effective multisensory integration in higher-order cortical areas. Such medium correlations indeed exist in the experimental data (Figure 2C). This result makes the prediction that alterations in the correlations in early spontaneous activity might affect later multisensory integration, which can be empirically tested. For instance, spontaneous activity in V1 has been shown to have different correlation properties in a mouse model of Fragile X [65] and in cases of disrupted inhibition [19, 47]. Our model therefore suggests that multisensory integration in RL may be perturbed in these cases. A key future test will be to perturb the temporal correlation structure of early V1/S1 spontaneous activity directly and determine whether this alters the alignment of V1/S1 maps in RL and the emergence of bimodal RL neurons.

What might be the origin of the correlated activity we observed in the visual and somatosensory cortex (Figure 2)? One possibility is that the source is external to the animal [66], i.e. a single stimulus with both visual and somatosensory aspects. Indeed, while a substantial portion of activity in the developing cortex occurs independently of sensory stimulation [23], both the immature visual [67] and somatosensory [68] cortices exhibit some degree of sensory-driven activity. Although it is unlikely that this external stimuli gives any topographic information before eye opening, it could help to correlate the activity between sensory cortices. Alternatively, subcortical areas like the (primary or higher-order) thalamus are in an ideal position for coordinating activation across distinct regions of the cortex [29, 56, 69]. The developing thalamus exhibits spontaneous propagating waves, which have been shown to guide the formation of sensory maps to the cortex and support circuit plasticity [56,70,71]. Spontaneous calcium waves propagate among the different sensory-modality thalamic nuclei up to the cortex and hence provide a means of communication among sensory systems [71]. Finally, correlated events in V1 and S1 could also originate from lateral long-range connections between the visual and somatosensory cortex. These types of connections emerge during the first two postnatal weeks in mice [72] and potentially directly propagate activation from one area to another [58, 59]. More specifically, it has been shown that postsynaptic excitatory neurons in a subregion near the anterior border of V1, situated in close proximity to S1, receive input from local fast spiking inhibitory neurons that have direct cortico-cortical input from the posterior area of S1 [73]. This cross modal suppression may contribute to these medium level correlations during development.

### Precise timing of map formation

We have shown that the somatosensory cortex’s early maturation is able to drive the aligned topographic development of the visual cortex Figure 4. This agrees with previous work showing earlier development of somatosensory cortex. For example, layer 4 (L4) to L2/3 excitatory synapses in somatosensory cortex were shown to mature 2 weeks earlier than those in visual cortex [74]. A similar template for the maturation of one map provided by another is observed in the refinement of cortico-collicular connections. Projections from S1 to the superior colliculus arrive as early as postnatal day 2 while projections from V1 enter the colliculus around postnatal day 6 [75], giving more time for S1 maps to refine. The precise timing of connectivity refinement between S1 and the superior colliculus likely guides the later developmental connectivity refinement between V1 and the colliculus and helps align the two maps [75, 76]. Moreover, this process of visual and somatosensory map alignment in the superior colliculus seems to be activity-dependent, with spontaneous activity in the retina playing an important role by driving the alignment of cortico-collicular connections from V1 to a genetically-induced duplicated retino-collicular map [76]. This modality difference in map refinement with the somatosensory leading the visual map seems to be present also in the refinement of connections between S1 vs. V1 and RL as we have shown earlier. Although the activity-dependent plasticity mechanisms behind this are experimentally unknown, inspired by earlier works [28, 52], our models suggest that similar activity-dependent plasticity involving Hebbian and heterosynaptic terms are sufficient.

### Role of recurrent connectivity

Neural dynamics in the adult cortex are strongly affected by recurrent connections [77, 78]. In our model, we have chosen to focus on feedforward connections between the primary sensory and RL for several reasons. First, recent experimental evidence from the developing somatosensory cortex suggests that feedforward connectivity might be established first [79], and second, the strength and functional specificity of recurrent connections in the vsual cortex increases only later in development [80]. Furthermore, abstraction is important for deriving robust biological insights from models [81] and here we are focused on how these feedforward connections arise and refine themselves over development building on our previous work of refinements between the thalamus and V1 [28].

### Uni- and bimodal neurons in other species

Neurons responding to only a single or to multiple modalities are common in other animals apart from mice, like macaques [82–84], cats [85–87], or ferrets [88, 89]. The surprising presence of unimodal neurons in association cortices has inspired a number of studies attempting to explain their functional significance [87, 90, 91]. Close examination of the receptive fields of neurons in the associative cortex of the cat revealed that stimulus location is likely represented in a distributed population code and that unimodal neurons could contribute by improving the accuracy of the encoded stimulus location [87]. Furthermore, the response patterns of unimodal neurons in the cat extrastriate cortex revealed that while these neurons do not respond to their non-preferred modality, their activation is nonetheless modulated when non-preferred and preferred stimulus occur at the same time [90], further suggesting that unimodal neurons contribute to a population code. Varying the proportion of uni- and bimodal neurons might thus contribute to a continuum of multisensory responses across different cortical areas [92]. Thus, our proposed explanation for the mixture of uni- and bimodal neurons in terms of optimal decoding of input origin integrates well into the existing literature. A direct experimental test of this decoding prediction will require simultaneous recordings from V1, S1, and RL at sufficient spatial resolution. The two-photon recordings used here were acquired sequentially across areas, and the wide-field recordings, while simultaneous, do not provide cellular-resolution activity in RL. Future simultaneous multi-area recordings could therefore test whether the ability to reconstruct V1 and S1 activity from RL activity increases across development, as predicted by the model.

Together, our results suggest that structured spontaneous activity can provide a developmental scaffold for the emergence of aligned multisensory maps and efficient multimodal representations in higher-order cortex.

## Methods

### Experimental procedure

#### Animals

All experimental procedures were approved by the institutional animal care and use committee of the Royal Netherlands Academy of Sciences. C57BL/6J mice of both sexes were used. All animals were aged between postnatal days (PN) 8-16. Mice of this strain open their eyes at P14.

#### *In utero* electroporation

For wide-field imaging (Figure 2) pyramidal neurons in layer 2/3 of the visual cortex were transfected with GCaMP6s (2 mg/ml) and DsRed (2 mg/ml) at embryonic day (E) 16.5 using *in utero* electroporation [93]. Pregnant mice were anesthetized with isoflurane and a small incision (1.5–2 cm) was made in the abdominal wall. The uterine horns were carefully removed from the abdomen, and DNA was injected into the lateral ventricle of embryos using a sharp glass electrode. Voltage pulses (five square wave pulses, 30 V, 50-ms duration, 950-ms interval, custom-built electroporator) were delivered across the brain with tweezer electrodes covered in conductive gel. Embryos were rinsed with warm saline solution and returned to the abdomen, after which time the muscle and skin were sutured.

#### Surgery

Animals were anesthetized with isoflurane (3 percent in 1 l/min O_2_). After anesthesia had become effective, lidocaine was used for local analgesia and a head bar with an opening (Ø 4 mm) above the visual cortex (0.5–2.5 mm rostral from lambda and 1–3 mm lateral from the midline) was attached to the skull with superglue and dental cement. For calcium imaging, a small craniotomy above the visual cortex was performed. The exposed cortical surface was kept moist with cortex buffer (125 mM NaCl, 5 mM KCl, 10 mM glucose, 10 mM HEPES, 2 mM MgSO_4_ and 2 mM CaCl_2_ [pH 7.4]).

#### Calcium indicator application

For the imaging experiments shown in Figure 1, the calcium-sensitive dye Cal-590 (Aat Bioquest) was dissolved in 4 *µ*l pluronic F-127, 20 percent solution in DMSO (Invitrogen) and further diluted (1:10) in dye buffer (150 mM NaCl, 2.5 mM KCl, and 10 mM HEPES) to yield a final concentration of 0.5 mM. The dye was then pressure-ejected into S1, RL and V1 at 10–12 psi for 8–13 min with a micropipette (3–5 MΩ) attached to a picospritzer (Toohey).

### Image acquisition

One hour before imaging, anesthesia was stopped, and animals were recorded in the absence of anesthesia. In vivo calcium imaging with cellular resolution (Figure 1) was performed on a Nikon multi-photon microscope (A1R-MP) with a 0.8/16x water-immersion objective and a Ti:Sapphire laser (Chameleon II, Coherent) Nikon software. Pixel size was 650 nm and images of 330 by 330 *µ*m were recorded at 5-10 Hz.

*In utero* electroporated pups were used for wide-field calcium imaging of visual and somatosensory cortex (Figure 2). Calcium events were recorded with a Movable Objective Microscope (MOM, Sutter Instrument). Time-lapse recordings were acquired with a 4x objective (0.8 NA, Olympus) and blue light excitation from a Xenon Arc lamp (Lambda LS, Sutter Instrument Company). A CCD camera (Evolution QEi, QImaging) was controlled by custom-made LabVIEW (National Instruments) based software and images were acquired at a frame rate of 20 Hz.

### Image processing

Images were processed as described previously [47]. For two-photon image processing, drift and movement artifacts were removed from each recording using NoRMCorre [94]. Each recording was aligned to the first recording in the series to remove any movements between recording sessions. Delta F stacks were made using the mean fluorescence per pixel as baseline, F_0_. ROIs were hand-drawn using ImageJ (NIH). Automated transient detection and further data processing was performed using custom-made MATLAB software (MathWorks).

Epifluorescence image processing: Delta F stacks were made using the average fluorescence per pixel as baseline. V1 was identified based on activity coordinates and shape after this method of identification was confirmed through immunohistochemistry for vGluT2 [19]. Recordings were performed in an area clearly defined by lambda and the brain’s midline. Our experience shows that the field of view varies across experiments only by approximately 200 *µm*. To precisely overlay the field of view with the boundaries of sensory areas, we use the extent of individual network events and the organization of our topographic maps that helped identify these boundaries, in particular those of V1. We used previously published cortical area outlines as a reference map [9]). Supplementary Figure S3 shows examples of how activity patterns and topographic organization reveal the outlines of sensory cortical areas.

### Statistical analysis

In two-photon recordings (Figure 1), activations within each cell were detected as in [47] as a peak with both an absolute height and relative prominence of at least 5% ΔF/F_0_. Participation rate was determined by % of active cells during an event out of total cells for that region. When more than 5% of recorded cells were active simultaneously, the activation was considered an event. Event duration was calculated as length of time above half max of the peak of activation. For previously published analyses of the wide-field calcium imaging data [47] (Figure 2), events were detected as calcium peaks with an amplitude above 2x the standard deviation. However, the functional correlation maps and lagged cross-correlation analyses shown in Figure 2 were computed from continuous fluorescence time courses and did not require event detection.

### Functional correlation maps

Functional correlation maps were generated as in [29], from three seed areas, each 10 × 10 pixels. For the main analysis in Figure 2, seed areas were placed in RL to determine whether distinct RL locations were correlated with spatially distinct regions in V1 and S1. For the additional examples in Supplementary Figures S1 and S3, seed areas were placed in V1, S1, or RL, as indicated in each panel. For each of the three chosen seed areas, the time course of each seed area was computed by averaging pixel values within the seed region for each frame. We then computed the Pearson correlation coefficient between each seed time course and the time course of every pixel in the field of view. The three resulting correlation maps were combined into an RGB image, with each color channel corresponding to one seed location. In Figure 2E and the top rows of Supplementary Figure S1, the RGB values represent the raw correlation values for the three seed locations. In Figure 2F and the bottom rows of Supplementary Figure S1, each pixel is assigned to the color channel with the highest correlation coefficient, and pixels below the chosen correlation threshold are shown in black. Supplementary Figure S3 uses the same functional-correlation-map approach to illustrate how cortical-area outlines were assigned.

### Linear mixed model

To correctly account for the nested data structure commonly encountered in neuroscience research [45], we fitted a linear mixed-effects model that treats animal and sample identity as random factors. We included interaction effects between age and region within our fixed effects. No transformations of the data were performed. Estimated effect sizes are shown in Figure 1H and Supplementary Tables S1-S3. Effect sizes on cortical region affected the intercept, but age and age-region interaction estimates affected the slope of the regression line. One-hot encoding was used because the regions are categorical variables. This means that one region, i.e. V1, was chosen as baseline and every other region was compared to this baseline. Any differences due to region were added/subtracted from the intercept for V1. Age and age-region interaction effects changed the slope and were therefore added or subtracted from the estimated slope for region V1. For Figure 1H, V1 was used as the reference region. To provide direct comparisons between all cortical areas, we also refit the same model using S1 or RL as the reference region; these results are reported in Supplementary Tables S1 and S3. Since PN8 was the earliest age used in our data, 8 was subtracted from all ages so that PN8 is the estimated intercept. Our linear model could not be valid outside of the ages used (PN8-PN16) and no extrapolation should be done without experimental confirmation. This multilevel analysis allowed us to examine the effects of age and cortical region on four response variables: amplitude, event rate, participation rate, and duration. To obtain p-values, we used Satterthwaite’s degrees of freedom method, as implemented in the *lmerTest* package [95] in R. The linear mixed model formula used was:

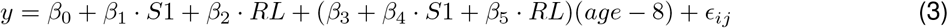

where *y* is one of the response variables, i.e. amplitude, event rate, participation rate, or duration and age is postnatal day. *β*_0_ is the intercept for V1 at age PN8, and *β*_1_ and *β*_2_ are the differences in intercept for cortical region (S1 and RL are dummy variables set to 0 if not predicting that region or 1 if predicting that region). *β*_3_ is the slope for V1 over age and *β*_4_ and *β*_5_ are the differences in slope (age-region interaction effects) from V1 over age for the cortical region of interest. *ϵ*_*ij*_ is the error from both random effects (mouse ID and sample ID) and residual error. When S1 or RL was used as the reference region, the same formula and age-centering were used, but the dummy variables were redefined relative to the corresponding reference area.

### Rate-based network model

Extending the model introduced by [28], we studied a feedforward, rate-based network with two one-dimensional input layers, **v** and **s**, and one output layer **r** with *N* ^*v*^, *N* ^*s*^, and *N* ^*r*^ neurons each. We used periodic boundary conditions in each layer to avoid edge effects. The initial connectivity between *v* and *r, W*_*v*_, and between *s* and *r, W*_*s*_, was initially all-to-all with uniformly distributed weights in the range *w*_ini_ ∈ [*a, b*]. In addition, a topographic bias was introduced by modifying the initially random connectivity matrix to have the strongest connections between neurons at the matched topographic location. These biased weights decayed with a Gaussian profile with increasing distance from the matched topographic location (Figure 3), with amplitude *a* and spread *s*. Note that we used the subscripts ‘s1’ and ‘v1’ to denote which brain area the parameters referred to. In simulations in which we varied the initial S1-to-RL connectivity bias, we varied the amplitude *a*_*s*1_ of this Gaussian bias while keeping its spread fixed. During the evolution of the weights, soft bounds were applied on the interval [0, *W*_max_].

The input populations *v* and *s* received spontaneous activation from a homogeneous Poisson process with an inter-event interval of 150 ms, implemented using a Binomial random variable with sufficiently small time steps of 1 ms. S1 and V1 neurons were driven by a combination of independent and shared spontaneous events. Each independent event was generated separately in either V1 or S1 and activated a randomly chosen, contiguous set of neurons. The percentage of neurons activated during an event was determined by 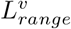 or 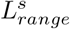 which was randomly chosen from a uniform distribution between 20% and 80% (See Table 1). The activated neurons were topographically next to each other with periodic boundary conditions. A shared event in V1 and S1 activated the identical set of contiguous neurons, topographically matched between V1 and S1. When shared events were included in the simulation, the inter-event interval in each input area was kept at 150 ms by splitting the total event rate into one shared Poisson process and two area-specific independent Poisson processes. If *p*_shared_ denotes the fraction of shared events and *λ* = 1*/*150 ms^−1^ is the total event rate per area, shared events were generated with rate *p*_shared_*λ*, and independent events in each area were generated with rate (1 − *p*_shared_)*λ*. Thus, the total event rate experienced by each input area was *p*_shared_*λ* + (1 − *p*_shared_)*λ* = *λ*, while the fraction of events shared between V1 and S1 was varied. A spontaneous event increased the firing rate of a fraction of neighboring neurons to *L*_amp_ for *L*_*dur*_ seconds. The activity of the output layer **r** was given as a leaky integrator of the input populations,

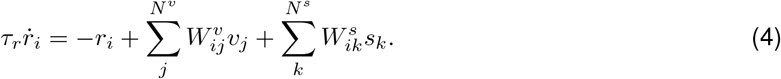

We studied the weight evolution under activity-dependent learning rules. In particular, following [28] we chose a rule that combines Hebbian and heterosynaptic effects that can produce selectivity,

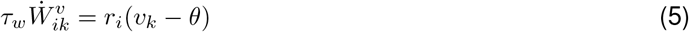

and

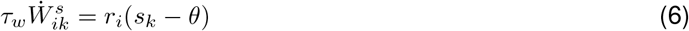

where *θ* is a heterosynaptic offset constant that defines a distance-dependent heterosynaptic depression. Heterosynaptic plasticity occurred when a nearby unstimulated synapse stabilized or depressed due to the plasticity in a stimulated one. This helped to stabilize the Hebbian plasticity [52]. Table 1 lists all parameters. For simulations with varying correlation values (Figure 3), the percentage of correlated events was chosen from a uniform distribution between 0 and 1, U[0,1]. For simulations with varying S1 bias, the Gaussian spread was fixed at 4, and the amplitude of the S1-to-RL Gaussian connectivity bias, *a*_*s*1_, was chosen to be 0.2, 0.4 or 0.6 (Figure 4A,B), or chosen from a uniform distribution between 0.05 and 0.4 (Figure 4C-E). Additional simulations were performed to test whether S1-like activity statistics alone could account for map alignment. In these simulations, *a*_*s*1_ was varied between 0.05 and 0.5, while S1 inter-event interval was varied between 130 and 150 ms or S1 event amplitude was varied between 0.9 and 1 (Figure S2). Simulation time of 500 s was used.

**Table 1.** Parameters used for simulations.

| parameter name | value | description |
| --- | --- | --- |
| $N^v, N^s, N^r$ | 50 | Number of neurons |
| $w_{ini}$ | U[0.35,0.55] | Range of initial weights (U: uniform dist.) |
| $a_{v1}, s_{v1}$ | 0.05, 4 | Amplitude and spread of initial Gaussian bias for V1-to-RL connectivity |
| $a_{s1}, s_{s1}$ | 0.05-0.6, 4 | Amplitude and spread of initial Gaussian bias for S1-to-RL connectivity |
| $\tau_w$ | 50 | Weight-change time constant for Hebbian rule [s] |
| $\tau_r$ | 0.001 | Membrane time constant [s] |
| $W_{max}$ | 1 | Max weights |
| $\theta$ | 0.5 | Heterosynaptic offset constant |
| $L_p^v, L_p^s$ | 150 | Inter-event interval [ms] |
| $L_p^s$ | $\in [130, 150]$ | Inter-event interval [ms] |
| $L_{dur}^v, L_{dur}^s$ | 15 | Duration of event [ms] |
| $L_{range}^v, L_{range}^s$ | U[0.2,0.8] | fraction of neurons in the network activated during an event |
| $L_{amp}$ | 1 | Amplitude during event for baseline simulations |
| $L_{amp}$ | $\in [0.9, 1]$ | Amplitude during event in S1 in Supp. Fig. S2 |
| % correlated events | 50% | % of events correlated between V1 and S1 |

### Expected weight dynamics and bimodality threshold

Since the dynamics of the output layer change much faster than the weights, *τ*_*r*_ ≪ *τ*_*w*_, we made a steady state assumption as in our previous two-layer model [28] and computed

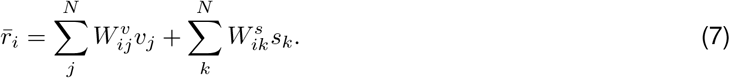

This allowed us to write the expected change in weight between (e.g.) the **v** and **r** populations as a function of the input statistics of the *v* and *s* population,

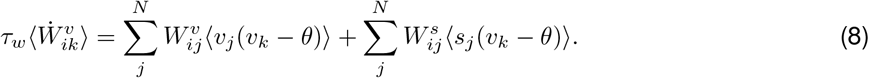

Here, the expectation is taken over the input activity statistics for a fixed connectivity state. Because of the separation of timescales between activity and learning, the weights *W*_*ij*_ are treated as fixed quantities during this calculation and can therefore be taken outside the expectation. This formulation allowed us to investigate the critical required amount of correlation between cells in **v** and **s** that permits the formation of stable bimodal cells in **r**. Specifically, we asked under which input correlation a depressed V1-to-RL synapse would become potentiated when the corresponding S1-to-RL synapse was already potentiated. In particular, we investigated the expected change of weight in the configuration where all connections between **v** and **r** were fully depressed and only one connection between **s** and **r** was potentiated:

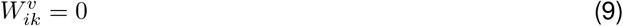

and

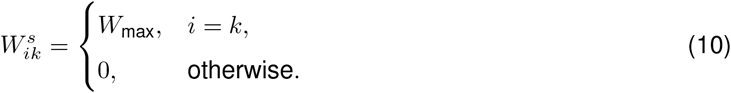

The expected weight change in this configuration was

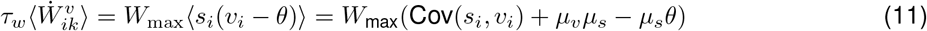

where Cov is the covariance and *µ*_*v*_, *µ*_*s*_ are the average firing rates. The critical amount of correlation, where potentiation and depression balance each other out, is characterized by the condition

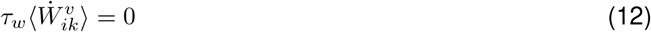

which we solved for the correlation, *γ*_*sv*_ = Cov(*s*_*i*_, *v*_*i*_)*/*(*σ*_*v*_*σ*_*s*_), as

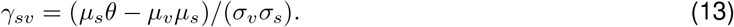

Assuming that rates and variances in **v** and **s** are equal and that the activity is Poisson, *µ* = *σ*^2^, the condition for the critical correlation further simplifies to

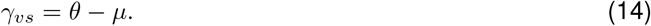

### Optimal weight matrix for correlated input populations

We considered the concatenated input activity at each time point, **X** = [**v**; **s**], of size *M* × 1, where *M* = *N* ^*v*^ + *N* ^*s*^ input neurons. This input drives activation in the “output” population, **r** = *ϕ*(**WX**), where *ϕ* is a (possibly non-) linear transfer function, **r** has size *N* ^*r*^ × 1 and **W** has size *N* ^*r*^ × *M*. The decoding problem introduced in Figure 5 used a sigmoid transfer function, i.e.

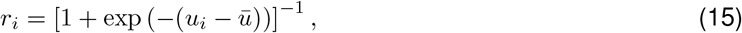

where *u*_*i*_ = (**WX**)_*i*_ and 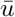 denotes the mean preactivation across RL neurons. However, to derive the optimal weight matrix analytically, a linear transfer function was used in order to show an analytical solution for this simplified case. This decoding problem could be phrased as inferring activity of the input populations from the output population, 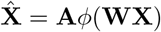, where **A** is the pseudoinverse of **W, A** = (**W**^*T*^ **W**)^−1^**W**^*T*^. We then searched for the optimal weight matrix, **W**_*opt*_ = [**W**^*v*^, **W**^*s*^] of size *N* ^*r*^ × *M*, where **W**^*v*^ represents the weights from **v** to **r**, and **W**^*s*^ represents the weights from **s** to **r**, that minimizes the squared error between **X** and 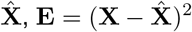 and therefore maximizes *R*^2^.

As the dimension of **r** is substantially smaller than the dimension of **X**, dim(**r**) ≪ dim(**X**), the optimal weight matrix needs to preserve the largest possible amount of variability of the input while projecting it into a lower dimensional space. Using insights from principal component analysis theory [96], we could demonstrate that the optimal weight matrix is composed of eigenvectors of the covariance matrix, ⟨**XX**^*T*^ ⟩, with the largest eigenvalues,

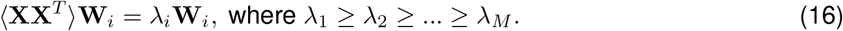

To better understand the structure of the eigenvectors, we could exploit the fact that in our set-up the covariance matrix, *C* = ⟨**XX**^*T*^ ⟩, is circular, i.e. every row of *C* can be obtained by a cyclic shift of the generating vector, *c* = [*c*_0_, *c*_1_, …, *c*_*M*−1_]:

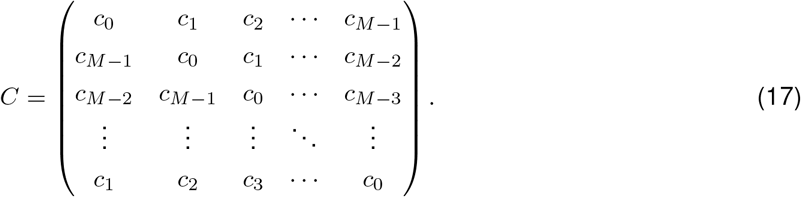

Eigenvectors and eigenvalues of circulant matrices have well-known closed form solutions [97]. The eigenvectors are the Fourier modes:

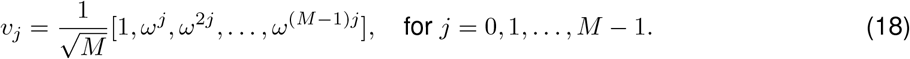

where *ω* = *e*^2*πi/M*^, a primitive *M*-th root of unity. The corresponding eigenvalues can be computed as

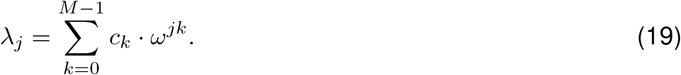

Since the covariance matrix is symmetric, i.e. *c*_*j*_ = *c*_*M*−*j*_ for all *j* and all values are real in the covariance matrix we rewrote the eigenvalues in terms of their real parts:

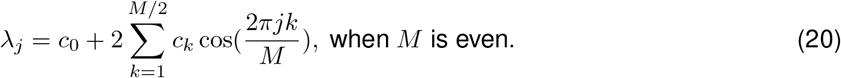

For a real-symmetric circulant matrix, the real and imaginary parts of the eigenvectors are themselves eigenvectors:

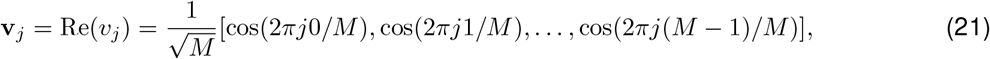

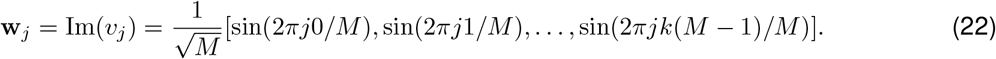

The first *M/*2 eigenvectors with nonzero eigenvalues from **v** and **w** will comprise **W**_*opt*_ which maximizes *R*^2^. Only the first *M/*2 are used since the dimension of **r** is *M/*2 in our case where *N* ^*v*^ = *N* ^*s*^ and can have a maximum of *M/*2 eigenvectors.

Our experimental set-up with correlated input populations translates into a generating vector *c* that has nonzero terms around the diagonal and offset from the diagonal by *M/*2. If the inputs between V1 and S1 are fully correlated, then *c*_*k*_ = 1 for *k* ∈ {0, …, *W/*2, (*M* − 1)*/*2 − *W/*2, …, (*M* − 1)*/*2 + *W/*2, (*M* − 1) − *W/*2, …, *M* − 1} and *c*_*k*_ = 0 else, where *W* is the extent of spatial correlation of the neurons. Taking this into consideration and removing any terms that simplify to zero due to the periodicity of the cosine function gives us the eigenvalues:

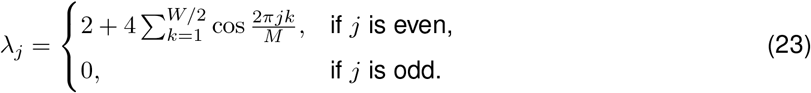

Hence, the eigenvectors with nonzero eigenvalue of the covariance matrix are exactly the first half of the eigenvectors of the sin and cos functions with an even-numbered period. Due to this simplification of the odd *j*’s leading to *λ*_*j*_ = 0 and even numbered periods, this results in eigenvectors that are identical on the indices {0, …, *M/*2 − 1} and {*M/*2, …, *M*} which is the definition of a bimodal cell and therefore all cells must be bimodal to maximize the *R*^2^ when input correlation = 1 between V1 and S1.

#### Topography

Network topography was defined to measure the degree to which the initial topographical bias is preserved in the final receptive field. To calculate topography, we calculated the Pearson’s correlation coefficient, *r*, of the final receptive field with an ideal receptive field template (Figure 3). The ideal receptive field template that was used has weights of 1 with a Gaussian spread of 4 from center.

#### Alignment

Alignment was defined to measure the similarity between the final receptive fields of V1 and S1. To calculate alignment, we calculated the Pearson’s correlation coefficient, *r*, between the final receptive field of S1 and the final receptive field of V1.

#### Reconstruction

To calculate the reconstruction of V1 and S1 activity from RL in Figure 5, we fitted a linear model to predict the combined V1 and S1 activity patterns from RL activity using 1000 s of simulated activity. Duplicate RL activity patterns were removed, and 80% of the remaining data was used for fitting to avoid over-fitting. Then the coefficient of determination was calculated as

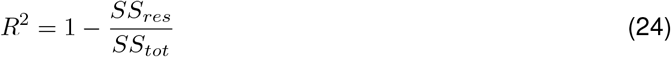

where *SS*_*res*_ is the sum of the residuals squared and *SS*_*tot*_ is the total sum of squares, using only one sensory area’s activity at a time, e.g. RL vs. V1 or RL vs. S1 activity compared with the linear fit of all 3 layers.

## Acknowledgments

We thank Mark Hü bener for feedback on the manuscript and all members of the Gjorgjieva and Lohmann labs for comments and discussions. This work was supported by the Max Planck Society (J.H.K., M.E.W., D.K. and J.G.), the Technical University of Munich (J.M.D. and J.G.), the European Research Council (ERC) under the European Union’s Horizon 2020 research and innovation program (Grant agreement No. 804824 to J.G.) and Nederlandse Organisatie voor Wetenschappelijk Onderzoek (grant numbers ALWOP.216, OCENW.KLEIN.535, 865.12.001, OCENW.M.22.310) to C.L., ZonMw (9126021) to C.L. and Stichting Vrienden van het Herseninstituut (805254845) to C.L.

## Author contributions

Conceptualization: J.G. and C.L. Experiments: N.Z., A.R., G.J.H. and P.P.M. Modeling and analysis: J.M.D., J.H.K., M.E.W., D.K. and J.G.; Writing: J.M.D., J.H.K., P.P.M., C.L. and J.G.

## Data and code availability

All code for generating the model and figures can be found at: https://github.com/comp-neural-circuits/MultisensoryIntegration.git

## A Supplementary Material

**Table S1.** Linear Mixed Model with V1 as reference. Values are estimates ± 95% CI width. Significance: * *p <* 0.05, ** *p <* 0.01, *** *p <* 0.001.

|  | Amps | Durations | Rates | Event Rate |
| --- | --- | --- | --- | --- |
| (Intercept) | $41.261 \pm 5.237^{***}$ | $7.524 \pm 1.685^{***}$ | $0.428 \pm 0.064^{***}$ | $1.377 \pm 0.755^{**}$ |
| Age | $-2.577 \pm 1.206^{**}$ | $-0.552 \pm 0.389^*$ | $-0.034 \pm 0.014^{**}$ | $0.334 \pm 0.167^{**}$ |
| Region S1 | $-8.533 \pm 1.557^{***}$ | $-0.778 \pm 0.467^{**}$ | $-0.008 \pm 0.044$ | $0.929 \pm 0.496^{***}$ |
| Region RL | $2.034 \pm 1.410^{**}$ | $-0.298 \pm 0.423$ | $0.025 \pm 0.040$ | $0.231 \pm 0.450$ |
| Age:Region S1 | $1.153 \pm 0.268^{***}$ | $0.058 \pm 0.080$ | $-0.011 \pm 0.008^{**}$ | $-0.176 \pm 0.085^{***}$ |
| Age:Region RL | $-0.317 \pm 0.241^{**}$ | $0.001 \pm 0.072$ | $-0.016 \pm 0.007^{***}$ | $-0.007 \pm 0.076$ |

**Table S2.** Linear Mixed Model with S1 as reference. Values are estimates ± 95% CI width. Significance: * *p <* 0.05, ** *p <* 0.01, *** *p <* 0.001.

|  | Amps | Durations | Rates | Event Rate |
| --- | --- | --- | --- | --- |
| (Intercept) | $32.717 \pm 5.239^{***}$ | $6.748 \pm 1.685^{***}$ | $0.420 \pm 0.064^{***}$ | $2.307 \pm 0.751^{***}$ |
| Age | $-1.423 \pm 1.209$ | $-0.494 \pm 0.389^*$ | $-0.045 \pm 0.014^{***}$ | $0.157 \pm 0.169$ |
| Region V1 | $8.510 \pm 1.558^{***}$ | $0.783 \pm 0.467^{**}$ | $0.010 \pm 0.044$ | $-0.929 \pm 0.496^{***}$ |
| Region RL | $10.555 \pm 1.336^{***}$ | $0.485 \pm 0.401^*$ | $0.034 \pm 0.038$ | $-0.698 \pm 0.422^{**}$ |
| Age:Region V1 | $-1.150 \pm 0.268^{***}$ | $-0.059 \pm 0.080$ | $0.011 \pm 0.008^{**}$ | $0.177 \pm 0.085^{***}$ |
| Age:Region RL | $-1.469 \pm 0.237^{***}$ | $-0.058 \pm 0.071$ | $-0.005 \pm 0.007$ | $0.170 \pm 0.075^{***}$ |

**Table S3.** Linear Mixed Model with RL as reference. Values are estimates ± 95% CI width. Significance: * *p <* 0.05, ** *p <* 0.01, *** *p <* 0.001.

|  | Amplitudes | Durations | Rates | Event Rate |
| --- | --- | --- | --- | --- |
| (Intercept) | $43.290 \pm 5.211^{***}$ | $7.225 \pm 1.678^{***}$ | $0.454 \pm 0.062^{***}$ | $1.609 \pm 0.732^{**}$ |
| Age | $-2.894 \pm 1.203^{**}$ | $-0.550 \pm 0.388^*$ | $-0.049 \pm 0.014^{***}$ | $0.326 \pm 0.165^{**}$ |
| Region V1 | $-2.045 \pm 1.411^{**}$ | $0.298 \pm 0.423$ | $-0.025 \pm 0.040$ | $-0.231 \pm 0.450$ |
| Region S1 | $-10.555 \pm 1.336^{***}$ | $-0.485 \pm 0.401^*$ | $-0.034 \pm 0.038$ | $0.698 \pm 0.422^{**}$ |
| Age:Region V1 | $0.319 \pm 0.241^{**}$ | $-0.001 \pm 0.072$ | $0.016 \pm 0.007^{***}$ | $0.007 \pm 0.076$ |
| Age:Region S1 | $1.469 \pm 0.237^{***}$ | $0.058 \pm 0.071$ | $0.005 \pm 0.007$ | $-0.170 \pm 0.075^{***}$ |

**Figure S1.**
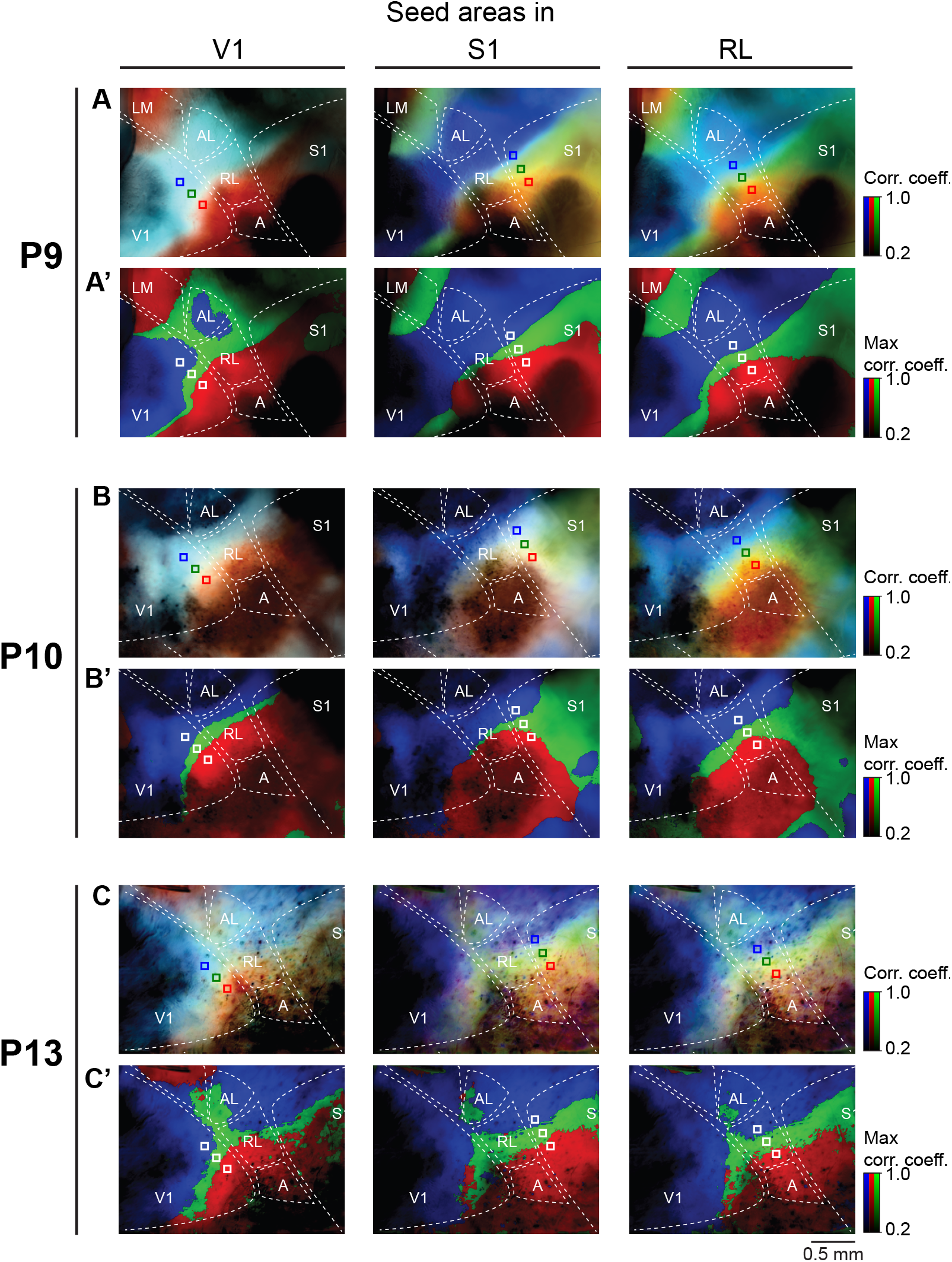
Functional correlation maps of 3 recordings at P9, P10 and P13. Pixel values represent Pearson correlation coefficients between each pixel and the three seed areas marked by blue, green, and red squares across the recording duration. Seed areas were placed in V1, S1, or RL, as indicated in each example. **A-C**. the color of each pixel is the RGB composite of the coefficients of the areas marked by the three colored squares. **A’-C’**. Same correlation values as in the top rows, but each pixel is assigned to the RGB channel with the highest correlation coefficient.

**Figure S2.**
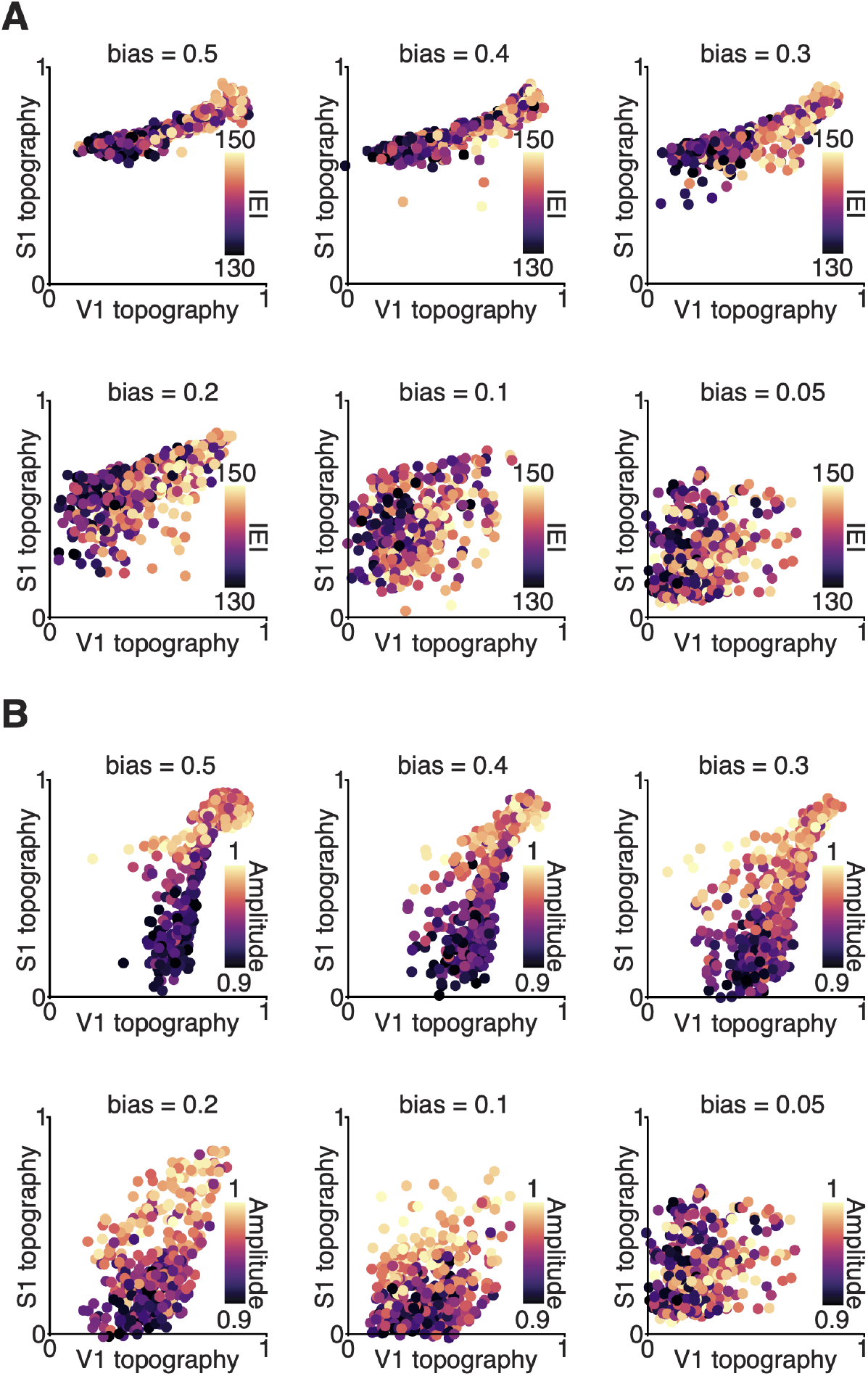
Effects of S1 activity statistics across different S1-to-RL connectivity biases. S1-to-RL connectivity bias was varied across simulations from 0.05 to 0.5. For all simulations, V1-S1 correlation was fixed at 0.5. **A**. V1-to-RL and S1-to-RL topography for different S1 inter-event intervals (IEI, in ms; indicated by color). **B**. V1-to-RL and S1-to-RL topography for different S1 event amplitudes (indicated by color).

**Figure S3.**
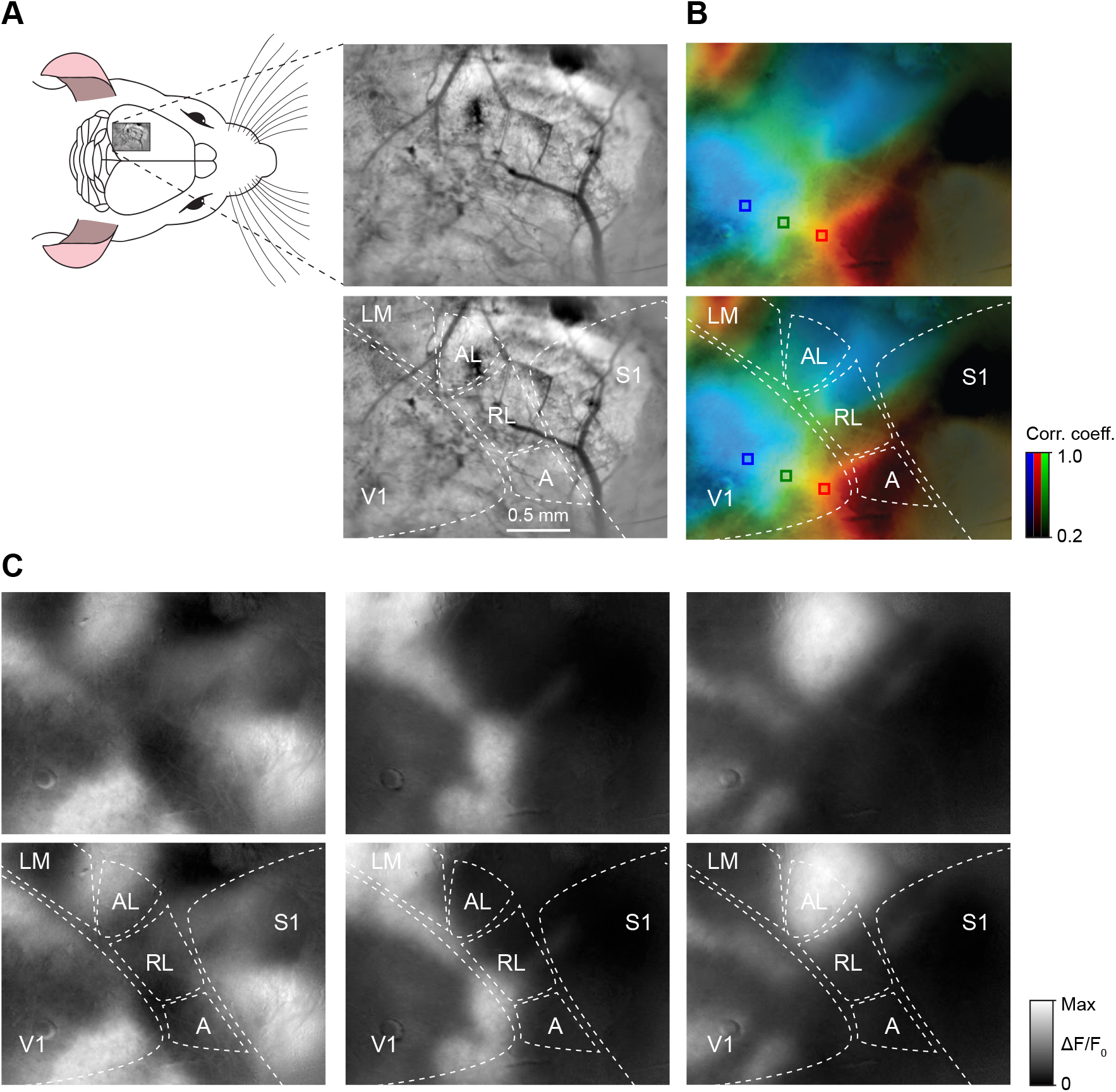
Mapping sensory cortex. **A**. Imaged cortical surface with sensory areas labeled (bottom). **B**. Functional correlation map, showing how correlated each pixel is with each of the seed areas shown as red, green and blue squares in V1 (for details see Supplementary Figure S1). Since topographic organization within V1 is monotonous, reversal of correlation distributions indicate areas outside V1, e.g., the red pixels in the top-left corner of the field of view. **C**. ΔF/F_0_ representations of individual network events in the imaged area. The extent of these events give a clear indication of the boundaries of V1.

